# Computing Signal Transduction in Signaling Networks modeled as Boolean Networks, Petri Nets, and Hypergraphs

**DOI:** 10.1101/272344

**Authors:** Luis Sordo Vieira, Paola Vera-Licona

## Abstract

Mathematical frameworks circumventing the need of mechanistic detail to build models of signal transduction networks include graphs, hypergraphs, Boolean Networks, and Petri Nets. Predicting how a signal transduces in a signaling network is essential to understand cellular functions and disease. Different formalisms exist to describe how a signal transduces in a given intracellular signaling network represented in the aforementioned modeling frameworks: elementary signaling modes, T-invariants, extreme pathway analysis, elementary flux modes, and simple paths. How do these formalisms compare?

We present an overview of how signal transduction networks have been modelled using graphs, hypergraphs, Boolean Networks, and Petri Nets in the literature. We provide a review of the different formalisms for capturing signal transduction in a given model of an intracellular signaling network. We also discuss the existing translations between the different modeling frameworks, and the relationships between their corresponding signal transduction representations that have been described in the literature. Furthermore, as a new formalism of signal transduction, we show how minimal functional routes proposed for signaling networks modeled as Boolean Networks can be captured by computing topological factories, a methodology found in the metabolic networks literature. We further show that in the case of signaling networks represented with an acyclic B-hypergraph structure, the definitions are equivalent. In signaling networks represented as directed graphs, it has been shown that computations of elementary modes via its incidence matrix correspond to computations of simple paths and feedback loops. We show that computing elementary modes based on the incidence matrix of a B-hypergraph fails to capture minimal functional routes.

## 1 Introduction

Cells must be able to receive, process, and respond appropriately to cues from their surrounding environment. The signal transduction component in the form of cellular elements that produce a response to cues from the cell’s environment will be referred from hereon in as a *signaling pathway*. The necessity of cells to process different signals causes several signaling pathways to interact with each other, creating signaling networks. The complexity innate to these networks, both from size and connectivity, makes computational modeling and analysis a requirement to understand how the cell communicates with its environment [30, 49]. It is well documented that malfunctions in signaling pathways from both epigenetic and genetic aberrations lead to several pathologies [42, 51], and malfunctions on signaling pathways, such as sustained proliferative signals, are a hallmark of cancer [35].

A signaling network is usually characterized as having a three-layer structure, with an input layer, an intermediate layer, and a target layer [103]. In mathematical models of signaling networks, the input layer nodes are typically ligands, exterior signals, receptors, or events that initialize the signal transduction process. The target layer, depending on level of abstraction, may be cellular responses, transcription factors, genes, metabolites, or processes that can be considered as the result of a complete signal propagation. The intermediate layer are the conduits of the signal, such as second messengers, kinases, and phosphatases. In Fig. 1 we show an example of a signaling network model for the Mitogen-activated protein kinase pathways originally introduced in [33]. Despite significant progress in understanding signaling networks, several kinetic parameters required for detailed mechanistic mathematical models remain elusive. Furthermore, certain signaling components, such as GTP-binding proteins, act as molecular switches, having an “active” or “inactive” status rather than a continuum. Thus, modelling frameworks circumventing the necessity of detailed kinetic parameters have been proposed in the literature, some of which, included in the present work, are reviewed by Samaga et al. [79]. Different formalisms can be translated, such as Boolean Networks to Petri Nets, and Petri Nets to a family of polynomials over a finite field [17, 86, 94]. This raises the question of how the formalisms used to capture signal transduction compare with each other.

**Fig. 1.**
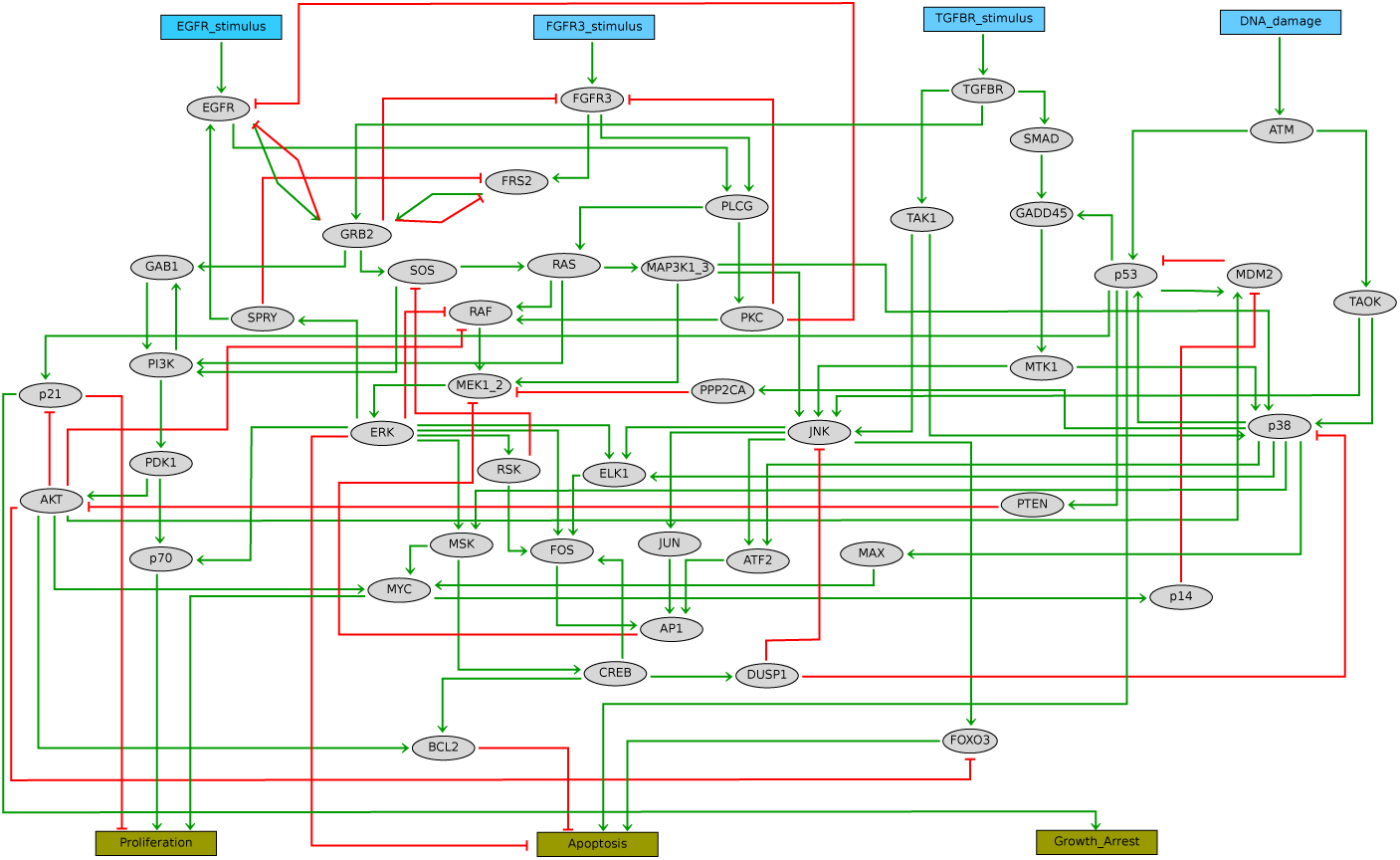
A model of the Mitogen-Activated Protein Kinase Pathway was downloaded from the GINsim model repository and adjusted using GINsim[16, 24]. The input layer (blue rectangles) is comprised of signals triggered by 4 different stimuli: (1) stimulus through the epidermal growth factor receptor (EGFR), (2) stimulus through the fibroblast growth factor receptor 3 (FGFR3), (3) stimulus of the transforming growth factor beta receptor I (TGFBR) and, (4) DNA damage. The output layer is comprised of Proliferation, Apoptosis, Growth Arrest. The intermediate layer (gray ellipses) is comprised of signal conduits, such as tumor supressors, kinases, transcription factors, and phosphatases. The arrow colors represent whether the action is inhibitory (red, blunt arrow) or activatory (green, arrowhead).

We provide an overview of arguably the more commonly used coarse-grained modeling frameworks of intracellular signaling networks: graphs, hypergraphs, Boolean Networks, and Petri Nets. We discuss the formalisms that modelers using these frameworks have proposed to capture how an intracellular signal transduces within a cell. We discuss the existing translations and describe different equivalences in the formalisms for capturing signal transduction. We adopt topological stoichiometric factories from metabolic network analysis tools [5, 1] and compare them to elementary signaling modes methodology [97].

To be sure, several criticisms and challenges exist for coarse-grained modeling frameworks, such as data discretization (see *e*.*g*. [55] for a comparison of time-series discretization methods), coarseness, and time-discreteness. At no point do we wish to argue that any of the modeling frameworks mentioned are the “be all end all” of modeling. What is worth remarking, however, is that in order to understand biology, modeling (widely defined), is necessary. To this date, no mathematical *model* is able to incorporate all the diversity of biological processes and spatial heterogeneity. Furthermore, biological data is intrinsically noisy [69]. Thus, detailed mechanistic models are often not feasible. As a result, a coarse-grained modeling approach is often necessary. A recent critical review [99] discusses in detail and provides references to the basis of using Boolean models in biology and also discusses some of the dynamics that cannot be captured by Boolean Networks, and many of the arguments there can be pushed to the frontiers of coarse-grained modeling frameworks.

## 2 Modeling frameworks for Signal Transduction Networks: an overview of their respective literature

We first introduce the modeling frameworks considered in this work and the mathematical formalisms necessary. We begin our discussion with perhaps the most fundamental representation of a signaling network; as a graph.

### 2.1 The basics of graphs

Let *G* be a directed graph *G* = (*V, E*), *V* = {ν _1_, …, ν _*m*_},, *E* ={*e*_1_, …, *e*_*n*_}. We say a node is a *source* (*sink*) node if it has no incoming (outgoing) edges. For example, in Fig. 1, Apoptosis is a sink node and EGFR stimulus is a source node. If all the nodes on a path *P*_*st*_ from *s* to *t* are distinct, we say *P*_*st*_ is a simple path. A *directed cycle* or *feedback loop* is a path from a vertex *s* to itself with no repeated vertices. Given an edge in a directed graph *e* = (*v*_1_, *v*_2_), *T* (*e*) = *v*_1_ and *H*(*e*) = *v*_2_ are the *tail* and the *head* of *e*, respectively. If *T* (*e*) = *H*(*e*), we will say *e* is a *self-loop*. The *incidence matrix* of a directed graph *G* = (*V, E*) is an *m*-by-*n* matrix 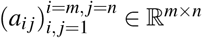

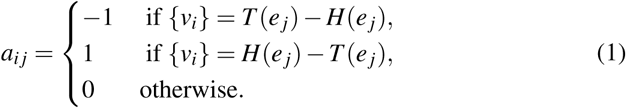

Notice that the incidence matrix does not represent self-loops.

A signed directed graph is a directed graph where each edge contains a positive (+) or negative (−) sign.

*Graph representations of signaling networks* Notice that the incidence matrix does not represent self-loops.

A signed directed graph is a directed graph where each edge is decorated by a + or − sign.

For intracellular signaling networks, its associated graph is commonly a signed directed graph hereon deemed *interaction graphs*, as in [45]. We note that the nomenclature *interaction graph*, is non-standardized. In the context of Boolean Networks, [48] use the nomenclature *interaction graph* for an unsigned directed graph. In interaction graphs, the nodes represent biological constituents (*e*.*g*. enzymes, metabolites, transcription factors), and edges represent interactions between the nodes. The signed edge stands for the type of interaction, *i*.*e*. activating (+) or inhibitory (−). For example, in Fig. 1, there is a positive (green) arrow from tumor protein p53 to mouse double minute 2 homolog (MDM2) gene representing the activation of MDM2 transcription by p53. The negative arrow from MDM2 to p53 represents the inhibition of p53 transcriptional activity, and subsequent degradation of p53 [96].

As previously remarked [45], it might not be possible to determine a direction of influence, or it might not be desirable to establish a direction of influence. Such interactions are best treated by different frameworks, or with a bidirectional edge [45]. If the interaction type is unknown, sometimes a label of 0 is used [103], although this is not a standard signed directed graph representation.

Weighted graphs have been used to represent protein interaction networks[84], where the weights stand for the reliability of the prediction of interaction of two proteins. Another graphical approach to represent signaling networks is to use bipartite graphs, such as *pathway graphs* [76]. A pathway graph is a bipartite graph *G* = (*M, I, E*) where *M* stands for the set of nodes of the network, *I* stands for the set of interaction nodes, and *E* is the set of edges connecting the nodes from *M* to nodes from *I* and vice versa. This methodology was used to model the epidermal growth factor receptor signaling network and is specially useful to analyze biochemical networks.

### 2.2 The basics of hypergraphs

Hypergraphs are generalizations of graphs, where the edges can connect more than two nodes. Analogously, we have directed hypergraphs.

A directed hypergraph is a pair 𝒢 = (𝒱, ℰ) where 𝒱 = (ν _1_,…, ν _*p*_) is a set of nodes and ℰ = {*e*_1_,…, *e*_*q*_} is a set of hyperedges. A hyperedge *e*_*i*_ is an ordered pair *e*_*i*_ = (*T* (*e*_*i*_), *H*(*e*_*i*_)) of nonempty subsets of 𝒱. *H*(*e*_*i*_) is the *head* of *e*_*i*_ and *T* (*e*_*i*_) is the *tail* of *e*_*i*_. If |*H*(*e*_*i*_) = 1 for all *i*, then the directed hypergraph is called a B-hypergraph [28]. Notice that when |*H*(*e*_*j*_) |= |*T* (*e* _*j*_) |= 1 for *j* = 1, …, *q*, the hypergraph is a standard directed graph (possibly with self loops).

As a word of caution, there are different definitions of hypergraphs in the literature. Ausiello et al. assume hyperedges are of the form (*S*, ν) where ν ∈ 𝒱 and *S* is a nonempty subset of 𝒱 [6]. Alternatively, Gallo et al. [28] assume that the head and the tail of a hyperdge are disjoint. The assumption that the tail and the head of a hyperedge are disjoint makes real biological processes, such as autocatalysis and self-regulators, difficult to model.

#### Definition 1

*Let 𝒢* = (𝒱, *ℰ*) *be a directed hypergraph. A* simple path *from s to t is a sequence of different nodes 𝒱*_0_ = (*u*_0_, …, *u*_*a*_) *and a sequence of different hyperedges E* = (*f*_1_, …, *f*_*a*_) *satisfying (1)-(3):*

1. *u*_0_ = *s, u*_*a*_ = *t*,
2. *u*_*i*_ ∈ *T* (*f*_*i*+1_) *for i* ∈ {0,…, *a*− 1} *and*,
3. *u*_*i*_ ∈ *H*(*f*_*i*_), *for i* ∈ {1, …, *a*}.

*We will say E* = (*f*_1_, …, *f*_*a*_) *is a* cycle *from s to t if t T ∈* (*f*_1_). *If a hypergraph has no cycles, we will say the hypergraph is* acyclic.

Given a directed hypergraph ℊ = (𝒱, ℰ) where 𝒱 = (*v*_1_,, *v*_*p*_), ℰ= *e*_1_,, *e*_*q*_, the *incidence matrix* of ℊ is a matrix whose rows correspond to the nodes and the columns correspond to the hyperedges. Namely, 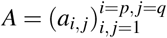 where

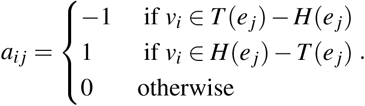

Notice that if one assumes that *T* (*e*_*j*_) ⋂ *H*(*e*_*j*_) = ∅ for all hyperedges, there is a one-to-one correspondence between directed hypergraphs and matrices with entries {0, 1,−1} through their incidence matrix [28].

We now discuss the concept of B-connection. Our definition of a B-connection comes from [72], although the concept of B-connection had previously been discussed [28].

Let 𝒢 = (𝒱, ℰ) be a directed hypergraph. For a node *u* ∈ 𝒱, let BS(*u*) = {*e*_*i*_ ∈ ℰ |*u* ∈*H*(*e*_*i*_)}. The definition of B-connection is a recursive one: given *s* ∈ 𝒱, we say *u* is B-connected to *s* ∈ 𝒢 if either *s* = *u* or there exists a hyperedge *e*_*i*_ ∈ BS(*u*) such that for each *w* ∈ *T* (*e*_*i*_), *w* is B-connected to *s*. The concept of *B*-connection is supposed to capture that all reactants must be present in order for a signal to be transduced [72]. The concept of B-relaxation distance has been recently introduced to provide a parametrization between hypergraph B-connectivity and graph connectivity [26]. Let ℬ _𝒢_ (*s*) be the set of hypernodes that are B-connected to *s* ∈ 𝒢. A B-hyperpath from *s* to *t* is a minimal subhypergraph Π_*s,t*_ such that *t* ∈Π_*s,t*_ is *B*-connected.

We remark that there is an alternative proposed definition definition of a B-hyperpath presented in [28], although a report of Nielsen and Petrolani (see the footnote in [90, p. 2595]) showed that the proposed definition is not equivalent to the definition of B-connection above if the directed hypergraph has cycles.

By the source layer of a hypergraph, we mean the set of source nodes in the hypergraph, *i*.*e*. a node with no incoming edge. We will assume hereinafter that our input layer is equal to the source layer. We will be using some concepts from metabolic network analysis, which we briefly introduce in the appendix, and refer the reader to [82, 5, 1] for the full treatment.

Hypergraphs remain underutilized in modeling for signaling networks [73], although their versatility of connecting more than two nodes at a time is advantegous for modeling reactions that produce more than one product or require more than one reactant [43, 73]. A hypergraph model of the WNT signaling network is presented in [72]. Hypergraphs can also be used to express logical relationships (AND, OR, NOT relationships) [45, 43], as done in [45] to model T-cell activation.

### 2.3 The basics of Boolean Networks

Boolean Networks in biology are usually attributed to the work of Kauffman [41] and Thomas [91].

#### Definition 2

*Consider a system X of n species (e*.*g. genes, proteins) to which we assign the variables x*_1_,…, *x*_*n*_. *Each of the variables takes a value in the set* {0, 1}. *A* Boolean Network *is a pair* (*k, F*) *where k* = {0, 1} *and F* = (*f*_1_,…, *f*_*n*_) : *k*^*n*^ →→ *k*^*n*^, *where each* local update function *f*_*i*_ : *k*^*n*^ → *k is a Boolean function in n variables*.

Different update strategies yield different dynamics, with subtleties: steady-state attractors are the same both for synchronous and asynchronous updates of a Boolean Network [19, 3]. Stable motifs [100] and trap spaces [46, 47], that is, trap sets with particularly simple dynamics, are also independent of the particular updating schedule strategy.

Boolean Networks have been widely used to analyze signaling networks. [78] constructed a 94-node and 123-interaction Boolean Network model of T-cell activation. [58] introduced a Boolean model to analyze pro-apoptotic pathways. A signaling pathway modeling the regulation of epithelial to mesenchymal transition (EMT) in primary liver cancer can be found in [88]. [104] provide a Boolean model for survival signaling in large granular lymphocyte leukemia. [13] constructed a 25 node Boolean Network incorporating the relationships between the NF-*κβ* pathway, a simplified apoptosis signaling pathway, and a necrosis signaling network, regulating how the cell “chooses” its fate. Several Boolean models of diverse systems, including intracellular signaling networks, are stored in repositories such as Cell Collective [37] and BioModels [20].

### 2.4 The basics of Petri Nets

We briefly introduce Petri Nets and refer the reader to [77] for details.

#### Definition 3

*Petri Nets are bipartite directed multigraphs, consisting of two types of nodes*, places *P* ={*p*_1_,…, *p*_*m*_} *and* transitions *T* = {*t*_1_, …, *t*_*n*_}, *and a set E of directed arcs weighted by natural numbers connecting only nodes of different types*.

*A place p* ∈*P in a Petri Net may carry any non-negative number of tokens m*(*p*), *called its marking*.

The dynamics of a Petri Net is given by its marking, and fireable transitions. Read-arcs are bidirectional edges representing requirement of the presence of markings but do not consume tokens. The incidence matrix of a Petri Net corresponds to the topology of the network [77]. Similar to self-loops in graphs and hypergraphs, the incidence matrix of a Petri Net does not represent read-arcs.

Chaouiya et al. and Steggles et al. [17, 86] describe translations from Boolean Networks to Petri Nets, depending on synchronous or asynchronous update rules. In particular, Petri Nets are able to display the dynamic properties of Boolean Networks, both under the synchronous and asyncrhronous updating strategies. Thus Petri Nets generalize the dynamics of Boolean Networks. Due to accumulation of the tokens, and consumption/production of tokens in the firing of transitions, Petri Nets can naturally describe types of biological processes such as biological consumption/reaction and inhibitions.

Introduced by Carl Petri [14], and later introduced in biology to analyze metabolic networks [71, 70], Petri Nets have been used to model diverse systems. A model of the pheromone response pathway in *Saccaromyces cerevisiae* appears in [77]. A model for the tumor necrosis factor receptor 1-mediated NF-*κ* B regulated signaling pathway appears in [4]. Li et al. [53] translate molecular interactions to Petri Net components, and used it to model an apoptosis network. [54] model a Petri Net as coupled “signal transduction components,” a set of substances that make an enzyme active. Coloured Petri Nets, where tokens are allowed to have different data types, are used by [103] to model signaling networks with only activations and reactions. The *Signaling Petri Net* [75], a synchronized Petri Net with an event generator, appeared in [67] to model the response of Langerhans cells to interferon regulatory factors. For a general overview of Petri Nets in biology, see [15, 50].

## 3 Capturing signal pathways in the different modeling frameworks

*Motivating question:* Given a set of source nodes 𝒳 and a set of target nodes 𝒥 in a signaling network, what nodes and edges are involved in transducing a signal from 𝒳 to 𝒥?

We seek to review the different forms of answering this question using the aforementioned formalisms.

### 3.1 In graphs

In graph models of signaling networks, signaling pathways are often represented as simple paths or shortest paths. For example, [45] compute feedback loops and simple paths from input nodes to output nodes from an interaction graph. Feedback loops in the topological representations of signaling networks are connected with the dynamics of the biological network [32, 12]. Paths and cycles are given an activating or inhibiting measure based on the parity of the sign. Klamt et al. [45] classify nodes as activators, inhibitors or ambivalent. Minimal path sets (MPS) [56] comprises of the set of all paths from input layer to target layer and feedback. *Sigflux* [56], a measure of importance of nodes in the network based on the amount of feedback loops and paths they are part of, uses the concept of MPSs. Lee et al. [52] proposed an algorithm for estimating how a signal propagates through a network purely based on an interaction graph. The algorithm predicts the direction of activity of nodes change in the network (down-regulated vs up-regulated) given an input. To compute all the nodes influenced by an input node, rooted trees have been used [103]. Similarly, reversing the directionality of the arrows, by computing rooted trees one might find the nodes which affect output nodes. Two-terminal series-parallel graphs were proposed as a method to capture parallel signaling pathways [84]. Nassiri et al. [62] weight the edges of the graph using the normalized similarity index [63]. Paths from input node to target nodes are weighted via a formula incorporating both node weights and edge weights, and the path with the highest weight is a likely candidate for a path from input to target node dominating the signaling process.

### 3.2 In hypergraphs

The concept of B-connection [72] describes the notion that all reactants must be present for a signaling reaction to occur. Constrain-based analysis on signaling networks, such as extreme pathways [80] and elementary flux modes [82] have been applied to signaling networks [64, 8]. Extreme pathways are special cases of elementary modes[65]. A similar definition to that of a B-hyperpath can be found in methodology for metabolic networks: the concept of a topological factory (appendix), which we adapt here since we will compare this to an object proposed in Boolean Networks. In the case of a metabolic network modeled as a hypergraph with stoichiometry matrix *S*, if *v* ∈ ℝ^| *ℰ* |^ denotes the flux of every reaction in the network (or in the case of quasi-stoichiometry, the amount of times a hyperedge is used), then *Sv* specifies the net production, or net change occurring. The notation (*Sv*)_*A*_ denotes the entries of *Sv* corresponding to *A* ⊆ 𝒱. Notice that this is equivalent to how the net change in the marking in Petri Nets is computed. The analogous concept of elementary flux modes [82] is the concept of a stoichiometric factory under the steady-state assumption. In B-hypergraphs, stoichiometric factories (see appendix) are unions of minimal topological factories [5].

### 3.3 In Boolean Networks

We now mention methodologies for analyzing signal transduction networks modeled as Boolean Networks. For a Boolean Network, we can apply techniques of analyzing signal transduction capabilities in directed graphs via its wiring diagram for computing signaling pathways. One advantage of Boolean Networks is the ability to exhibit dynamical information of the system in addition to their structural information. *Stable motifs* [100, 101] are a bridge between the structure and the dynamics of a Boolean Network. In particular, stable motifs are independent of the update scheduling strategy. Stable motifs are closely related to the concepts of trap spaces [47]. Using the MAPK pathway modeling cell fate decisions [33], the minimal trap spaces are computed and used to give a lower bound on the number of cyclic attractors [47]. To compute trap spaces, a graph expansion method is used; the prime implicant graph which is a *unique* hypergraph expansion of a Boolean Network not depending on any particular normal form.

Mashewari et al. [57] label the edges of the interaction graphs with causal logic, *e*.*g*. if 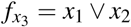, then the arrows from *x*_1_ to *x*_3_ and from *x*_2_ to *x*_3_ are labelled as sufficient arrows (permanent activation of either one of them suffices to activate *x*_3_ regardless of the rest of the network). These causal edges can sometimes be chained together to give information on the causal relationships of two nodes in the network that are far apart, and can be used to compute the logic backbone of the network, a vast simplification of the Boolean structure of the network. Furthermore, it can also be used to compute some stable motifs and for network reduction purposes [57].

An accepted technique to analyze Boolean Networks is based on the idea that long term behavior of networks is captured by their attractors [40]. Steady-state attractors are often associated with cellular phenotype and/or long-term cellular behavior [21, 3, 29]. Complex attractors (*e*.*g*. cycles), are often associated to oscillatory behaviors, such as progression through the cell cycle [3]. For example, some of the attractors in are associated to a proliferative phenotype in cancer cells [27]. Related to attractors, is the computations of the basin of attractors; the initial states that lead to an attractor. For attractor analysis, reduction techniques have been proposed in the literature; for example, using the concept of *Stable Motifs*[100], and via the polynomial dynamical system representation [93, 92]. Polynomial Dynamical Systems offer several advantages to logical representations of Boolean Networks, including the computation of steady states using computational algebra, and computational algebra tools for dimension reduction [95, 94, 93, 92]. Furthermore, they allow for a framework encompassing multi-state networks in which the tools of computational algebra allow for computation of steady states without the need to simulate the complete state space.

Due to the complexity of the state space (it grows exponentially with the number of nodes in the network), it is generally impossible to determine the complete dynamical evolution of a Boolean Network from its structure alone [39, 31]. Functional cycles, *i*.*e*. cycles that generate attractors, in the interaction graph of a Boolean Network are connected to the long term behavior of a Boolean Network [22]. For a survey of results connecting the dynamics and structure see [66].

An *Elementary Signaling Mode* (ESM) [97], based on structural analysis of a Boolean Network, is a minimal set of elements of the network that can perform signal transduction from initial node to nodes in the target layer. The network is expanded by introducing complementary nodes for nodes inhibited by other nodes or are inhibiting other nodes. Wang *et al*. [97] introduce a “composite” node to represent conditionally dependent relationships. See *e*.*g*. Fig. 2 and [97] for details.

**Fig. 2.**
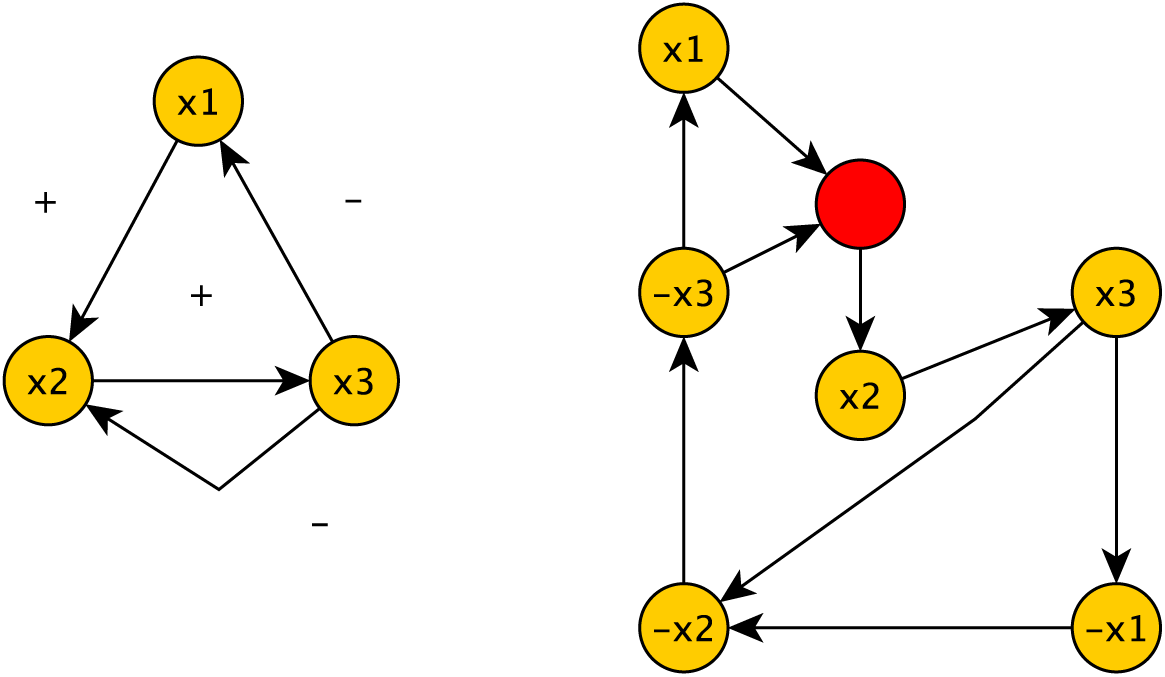
Consider the wiring diagram (left) of the Boolean Network 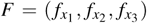, where 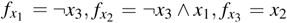 To expand the network, we add a complimentary node for each node of the network representing the absence or inactivity of a network component. These nodes are represented by −*x*_*i*_. The update functions for these nodes are the logical negations of the update rules of *x*_*i*_. For example, the update rule for −*x*_2_ is now −*x*_2_ = ¬*x*_1_ ∨ *x*_3_.

The expansion of a network provides a useful compromise between the wiring diagram (structure) and the full representation of a signaling network and rids the wiring diagram of some ambiguities. In fact, if the complete network expansion where both composite nodes and complementary nodes are added for every node in the network are used [2], the update rules can be read directly from the expanded network. For computations of elementary signaling modes, different signaling network expansions have been used such as expansions where composite nodes are added for logical dependencies[2], and complementary nodes are added for every node in the network. In contrast, other expansions only add composite nodes [89] to represent synergy. Therefore, one needs to first provide a mathematical definition of ESMs to make it computationally amenable in systems biology software. [98] provide a similar concept for graphs with dependent edges namely, the concept of a minimal functional route (MFR). Dependent edges represent necessary and sufficient conditions for signal transduction from a set of nodes *x*_1_,…, *x* _*j*_ to a node *y*. Again the network is expanded to a new network *Ĝ* by adding composite nodes to represent dependent edges (see Fig. 3 and refer to [98] for details).

**Fig. 3.**
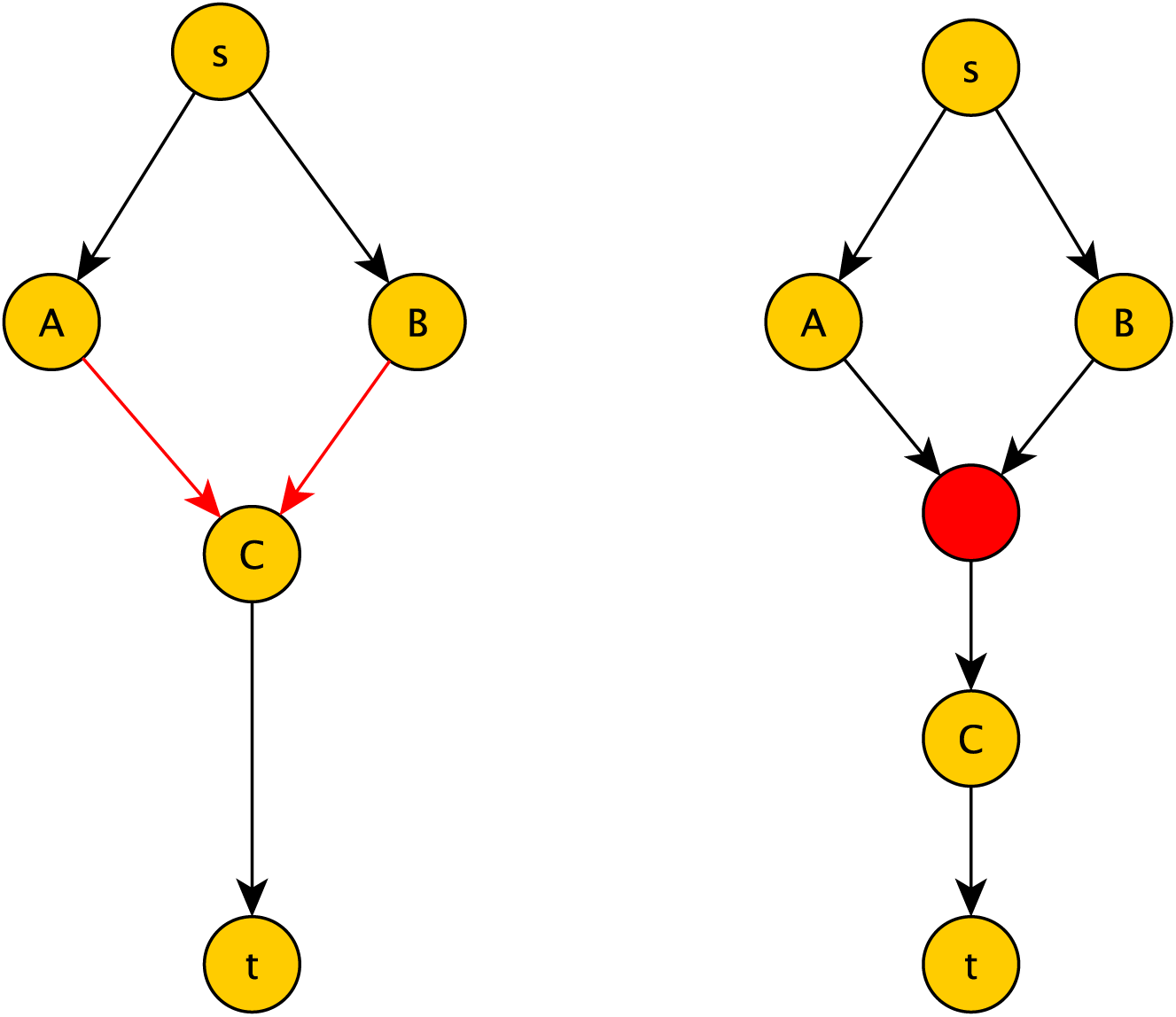
The edges in red are dependent edges. This represents that in order for node *C* to receive the signal, the signal from both *A* and *B* are necessary. On the right is the expanded graph, where a composite node is shown in red. In order for a signal to be transduced to *C*, both upstream nodes from the composite nodes must be included.

#### Definition 4

*(Minimal functional route (MFR) from [97]) Given a graph with dependent edges with source node s and sink node t, and its expanded graph Ĝ, an MFR from s to t in Ĝ is a minimal set of nodes and edges satisfying: s and t are in the set; each node is connected from s by simple paths; any original node other than the source node has one direct predecessor in the MFR, and any composite node has all of its direct predecessors in the MFR*.

In the special case of a graph with no dependent edges, computations of MFRs from *s* to *t* is the same as computing simple paths from *s* to *t* [98].

Computing simple paths or shortest paths from source node to target node might miss key information from the signal transduction process. For example, consider the graph on the left in Fig. 7. The only shortest path from *s* to *t* is the path *s, r*, 2, 3, where *r* is the red node. Therefore, computing simple paths would miss the fact that node 1 is necessary for a complete signal transduction process from *s* to *t*. In general, there is no relationship between the number of minimal functional routes and the number of simple (or shortest) paths from a source node *s* to a target node *t*, as we show in Fig. 4.

**Fig. 4.**
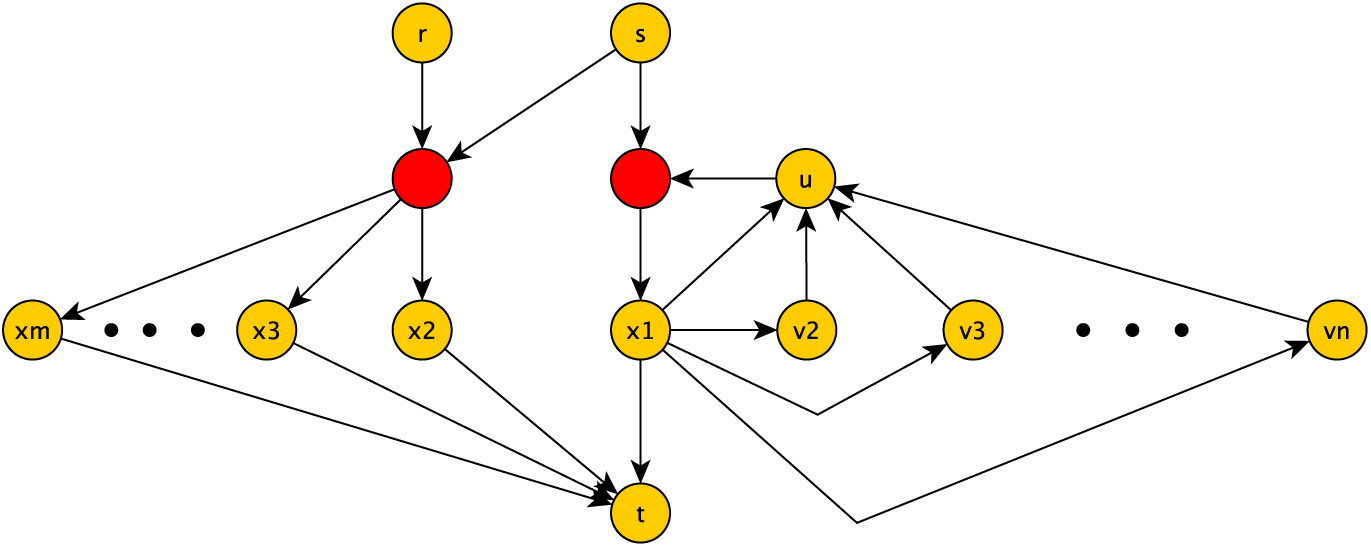
The composite nodes are shown in red. There are *m* simple (or shortest) paths from *s* to *t* and *n* minimal functional routes from *s* to *t*

### 3.4 In Petri Nets

Similar to Boolean Networks, Petri Nets allow modeling the dynamics of a signaling network. To analyze the long term behavior of a Petri Net, one is often interested in the behavioral properties of a Petri Net, such as computing the coverability graph of the Petri Net, the boundedness of the Petri Net and whether or not the net has dead markings [50]. Unfortunately, similarly to analyzing the state graph of a Boolean Network, many of these questions are computationally limiting, and thus one often resorts to structural analysis methods.

In the framework of Petri Nets, the commonly used methodology for capturing signaling pathways is T-invariants. In the case that there is a sequence of transitions realizing a vector **y**, a T-invariant **y** corresponds to a sequence of transitions that does not change the given marking [77]. In the framework of metabolic networks, minimal T-invariants are counterparts to elementary flux modes [102, 7], although elementary flux modes are more general due to the fact that reactions are allowed to be reversible. A place invariant is the counterpart of moiety conservation [7]. As previously mentioned, read-arcs are not reflected in the incidence matrix. Therefore, T-invariant analysis without taking into account this extra structure could yield T-invariants that are meaningless. To take this into consideration Sackmann et al. [77] introduce the concept of a feasible T-invariant to be a minimal set of transitions that can fire in sequence under the initial minimal marking, without changing the marking and discuss how to handle read-arcs for T-invariant analysis, such as considering read-arcs as unidirectional arrows or bridging T-invariants. The idea is that feasible T-invariants stand for minimal sub-entities of the Net relevant to capture signal transduction. The concept of feasible T-invariants is further developed to Manatee invariants [4], a minimal linear combination of T-invariants whose induced network is feasible under the initial marking.

## 4 Comparing different formalisms: A perspective

In this section we discuss some translations between the formalisms found in the literature. We also discuss what we see, from our perspective, as some of the advantages and disadvantages of the different formalisms.

An interaction graph is the easiest to construct, both from literature and data, and are easily interpretable. Signed directed graphs have been heavily studied by researchers above a broad spectrum of fields and efficient tools and algorithms have been developed to detect the signaling pathways in interactions graphs, such as shortest and simple paths, feedback loops and cycles. Several packages tools for network analysis are publicly available and user friendly, including Cytoscape [85], igraph [23], and NetworkX [34].

Once more information becomes available, either from the literature, or from experimental work, interaction graphs can be extended into a different framework. Hypergraphs allow to express co-operativity of components and allow a useful representation of Boolean Networks in the form of a logical hypergraph [45]. Some tools for the enalysis of hypergraph representations of signaling networks include the HALP Python library (available at https://tmmurali.github.io/halp), and the Bioconductor Hypergraph package [25].

Boolean Networks and Petri Nets each offer a dynamical model of a signaling network. Both Boolean Networks and Petri Nets have a large community of researchers in the biological sciences. Interestingly, every Boolean Network can be seen as a Petri Net [17, 86, 87]. Chaouiya et al. [17] give a translation from the Boolean framework to a 1-safe standard Petri Net (the multistate case can be found in [18]). Assuming one starts with a valid marking (the sum of the tokens between a gene and its complement is 1), the reachability graph of the corresponding Petri Net is equivalent to the fully asynchronous updating state graph of the corresponding Boolean Network. In particular, notice that given a valid marking, a realizable T-invariant is a multiset of transitions that does not change the given marking, and since the reachability graph and state graph of the Boolean regulatory network are equivalent, the counterparts of cycles in the state transition graph of a Boolean model are T-invariants. Stegless *et al*. [86, 87] provided a translation from Boolean Networks to Petri Nets focusing on gene regulatory networks for the synchronous updating timing schedule. A related approach can be found in [77], where insight on how to translate logical rules into the Petri Net framework is given. As previously remarked [50], Petri Nets are useful for representing consumption and production mechanisms, whereas Boolean Networks are more appropriate to model regulatory interactions (a regulator can alter the state of a target, whereas the state of the regulator does not change itself) [50].

The graph expansion method of [97] gives a hypergraph structure which can easily be translated into a Petri Net. However, the dynamics of the Boolean Network are not preserved under this translation. This expansion method is closely related to the translation from [17], although the latter heavily relies on either read-arcs or inhibitory arcs to preserve the dynamics. Similarly, the unique prime implicant graph of a Boolean Network is a *B*-hypergraph[46, 47], and can easily be translated into a Petri Net structure or into a graph with composite nodes. It should be further investigated how this translation preserves the dynamics.

From a modeling perspective, Boolean Networks are easier to set up than Petri Nets. Several tools exist to both model and analyze Boolean Networks [60, 16, 59, 95, 83, 11, 44].

Petri Nets are the most general, although they have a smaller track-record of being used in the modeling of intracellular signaling networks. However, Petri Nets allow the most flexibility in terms of the processes being modeled: they are natural representations of reaction networks, and due to read-arcs they allow flexibility in the signaling processes they can model. Interestingly, there are existing translations from 1-bounded Petri Nets to a family of Boolean Networks[94]. Under the given translation, the authors show that dead markings correspond to a solution of a system of polynomial equations over finite fields. Furthermore, recovering P-invariants and T-invariants of the original Petri Net is straightforward. For every place-transition Petri Net, there is an associated bipartite graph structure, and in particular, a hypergraph structure. In particular, we may apply the concepts for analyzing hypergraphs, such as hyperpaths and topological factories, to Petri Nets. Some tools to analyze Petri Nets include Snoopy [36], TINA [10], and LoLA [81], and the Integrated Net Aalyzer (INA) [38]. GINsim also includes methdos for the translation of logical models into Petri Nets Formats [17, 18, 16, 61].

Well-known reversible construction to create bipartite graphs from hypergraphs [43] exist. Naturally, every graph can be considered as a special case of a hypergraph. The signaling hypergraphs can be converted to standard graphs in two different ways [72].

We now further show how some methods in signaling networks are closely related to analogous constrained-based methods in metabolic networks.

### 4.1 Capturing signaling via topological factories

We show that minimal functional routes (MFRs), and thus elementary signaling modes (ESMs), are special cases of topological factories previously described [5, 1] once graphs with composite nodes are translated into a hypergraph structure. In particular, the concept of topological factories extends the concept of MFRs.

Let *G* be a graph with composite nodes [98] and no self-loops and with the added property that if (*x, c*_1_) is an edge where *c*_1_ is a composite node, then there is no edge (*c*_1_, *x*). Such graphs can be attained from the wiring diagram with no self-loops where we know which edges are dependent via the Boolean function of the nodes [97, 2, 89]. Furthermore such graphs can easily be converted to a B-hypergraph by collapsing incoming edges into a composite node *c*_1_ into the tail of a hyperedge (see Fig. 5). Due to the assumptions, we have a hypergraph with the property that for every edge *e* in the B-hypergraph, *H*(*e*) ∩*T* (*e*) = ∅.

**Fig. 5.**
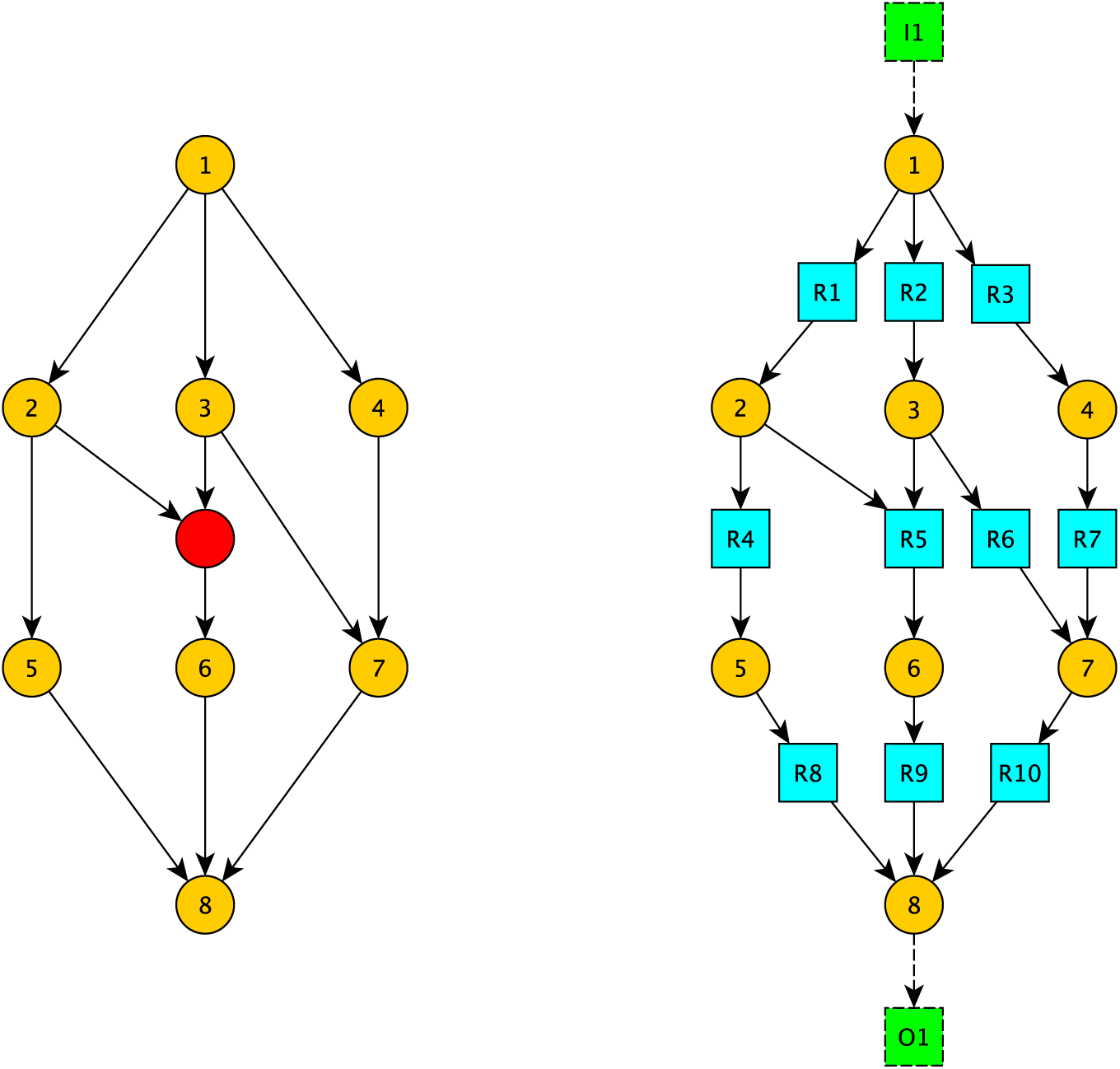
B-hypergraph from expanded graph. Adapted from [98, Figure 1(b)]. On the left, an expanded AND-OR graph. The composite node is shown in red, original nodes are shown in yellow. On the right, the corresponding hypergraph. The input node 1 has been connected to an environment transition *I*_1_ and the target node has been connected to an environment transition *O*_1_. for computations of elementary flux modes.

#### Theorem 1

*The set of minimal functional routes from s to a sink node t is contained in the set of minimal topological factories from s to t*.

#### Proof

This follows from the fact that every node in an MFR is connected to *s* by a simple path. That is, for every node *u* in the MFR distinct from *s*, there exists a hyperedge *e* in the MFR such that *H*(*e*) = *u*. Thus, if *E* ={*e*_1_,…, *e*_*m*_} are the hyperedges corresponding to the MFR, we have {*t*} ⋃*T* (*E*) ⊇*H*(*E*) ⋃{*s*}. Thus the hyperedges of the MFR form a topological factory from *s* to *t*.

Furthermore, if the expanded graph is an acyclic connected graph with a single source and a single target node, then computing the set of MFRs is the same as computing the minimal topological factories.

#### Theorem 2

*Let G* = (*V, E*) *be an acyclic connected graph with a single source node and composite nodes. Let t be a sink node where s* ≠ *t. Given a minimal topological factory U from s to t, the hypergraph induced by U is a minimal functional route from s to t*.

#### Proof

Since *G* is acyclic, there is an ordering of the nodes *w* : *V* → ℕ such that if (*v*_*i*_, *v* _*j*_) ∈*E*, then *w*(*v*_*i*_) *< w*(*v*_*j*_). Since *s* is the only source node, *w*(*s*) is the minimum value *w* attains on *V*. Since *U* is a minimal topological factory from *s* to *t, t* = *H*(*e*_1_) for some *e*_1_ ∈ *U*. Let *u*_1_ be any node in the tail of *e*_1_. Notice that *w*(*u*_1_) *< w*(*t*). If *u*_1_ = *s*, then there is a simple path from *s* to *t*. Otherwise, let *e*_2_ be the hyperedge containing *u*_1_ as its head, and let *u*_2_ be any node in the tail of *e*_2_. Proceeding in this way, we get a sequence of nodes *u*_*m*_, *…*, *u*_1_, *u*_0_ = *t* that are connected by a path with the property that *w*(*u*_*m*_) *< w*(*u*_*m*−1_) *<* …*< w*(*t*). Since *U* is a topological factory from *s* to *t*, this sequence must terminate at a node with no incoming hyperedge, *i*.*e. s*. Thus, we have created a simple path from *s* to *t*.

Now let *u* be any node different from *s, t* in the topological factory. Then *u* must be contained in the head of a hyperedge, and by minimality, a unique hyperedge. Applying the same method as above, we can create a simple path from *s* to *u*.

In expanded graphs with cycles, there are minimal topological factories from *s* to *t* that are not MFRs (see Fig. 6).

**Fig. 6.**
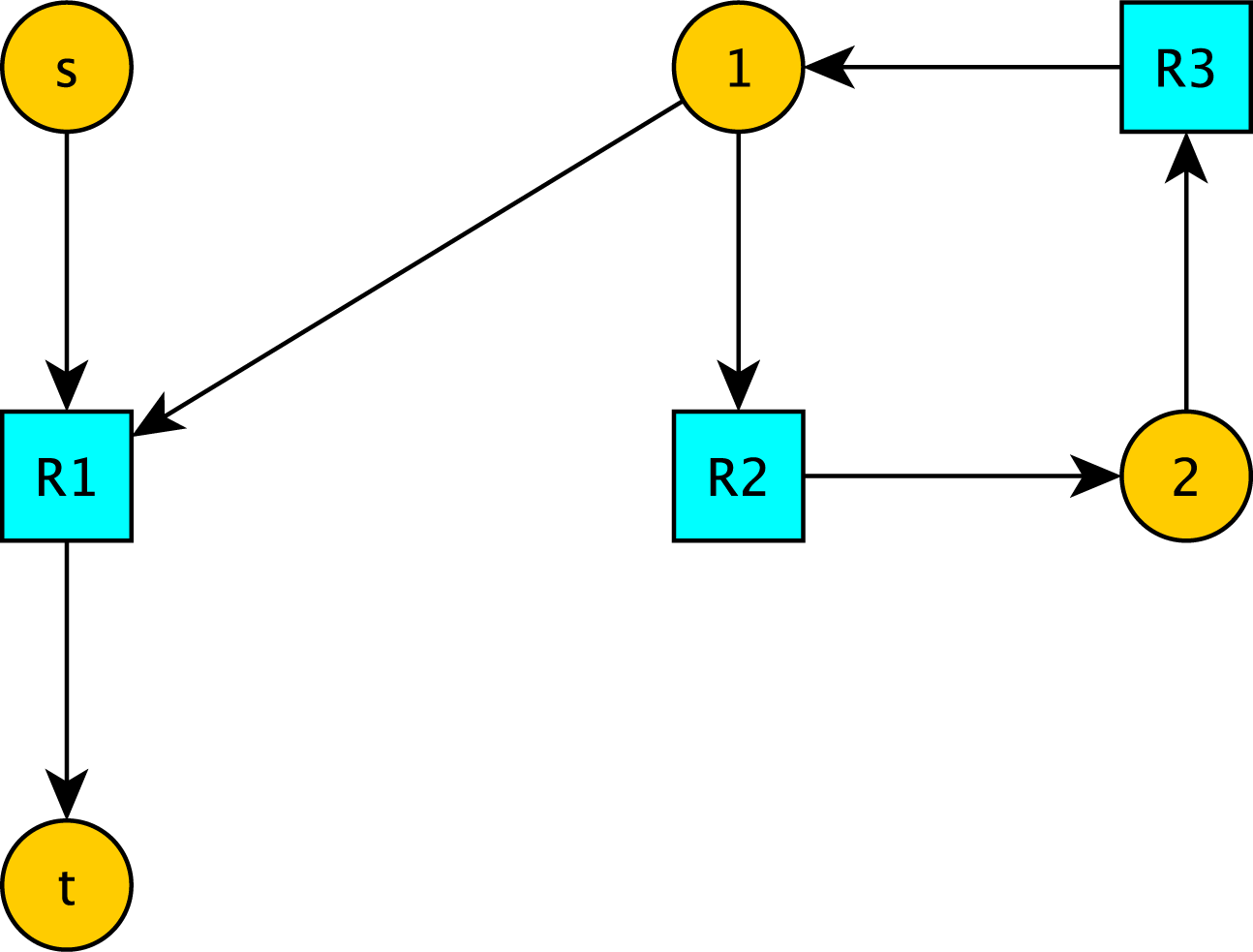
Not all topological factories are minimal functional routes from *s* to *t*. There is no MFR from *s* to *t* since transduction of a signal from *s* to *t* would require including node 1. However, there is no simple path from *s* to node 1. *R*_1_, *R*_2_, *R*_3_ is a topological factory from *s* to *t*. In the case the graph is acyclic, MFRs from *s* to *t* are the same as the topological factories from *s* to *t*.

**Fig. 7.**
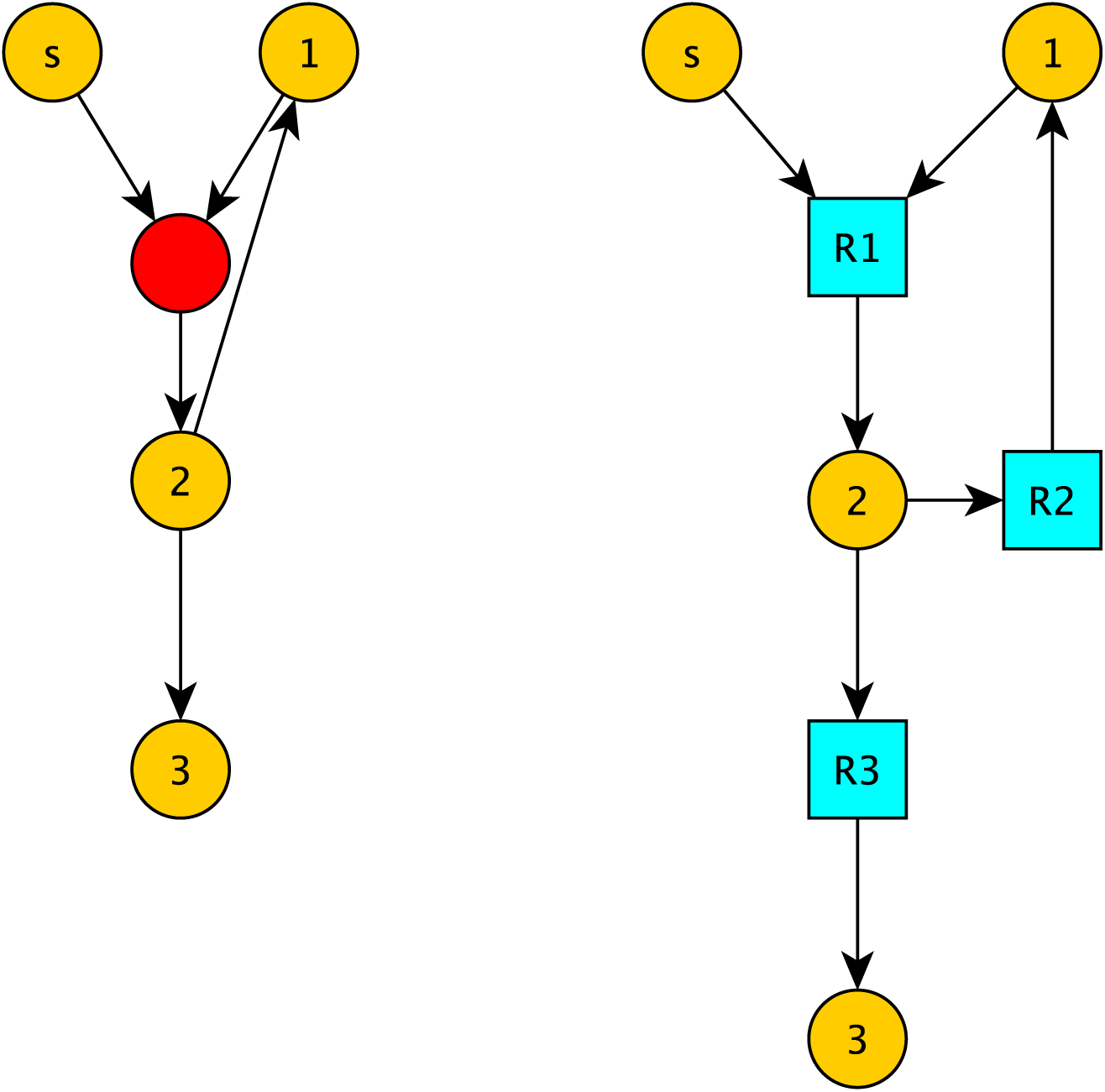
A minimal functional route (and thus topological factory) that is not an S-factory. Using the incidence matrix as the stoichiometric matrix for this hypergraph, there is no S-factory from *s* to *t*. However, it is a minimal functional route from *s* to 3.

Given a directed graph *G*, node *s* and node *t* and an incidence matrix *A*, there is a close relationship between simple paths from *s* to *t* and elementary modes of a slightly adjusted incidence matrix [45]. It is natural to wonder if for a given directed graph with dependent edges, the analogous process as in [45] can be used to compute MFRs via the incidence matrix of its respective hypergraph.

We will make no distinction between a mode and its representative *V*^*\**^ (see the Appendix, section 6.3 for the definition of modes). Notice that elementary modes, as formally defined in [82], are usually computed from the stoichiometry matrix of a metabolic network, and the restriction 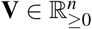 is not usually placed to allow reversible reactions.

Let ℋ be a directed hypergraph with no self-loops and let *A* be its incidence matrix. Let *M* be the set of MFRs from input node *I* to *O*. It is natural to wonder if computing elementary flux modes on *Ã*, where *Ã* is the matrix derived from *A* adding a column with a +1 in the row corresponding to *I* and a −1 in the row corresponding to *O* and zeroes everywherelse, will yield minimal functional routes and feedback loops.

We answer this question in the negative.

Consider the hypergraph corresponding to Fig. 7. The adjusted incidence matrix is as follows

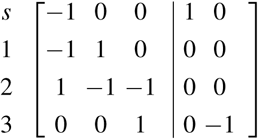

Which row reduces to:

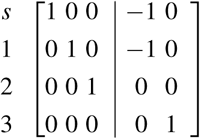

Thus the hyperarc *R*3 cannot be used, *i*.*e*. there is no elementary flux mode from *s* to 3. However, notice that the hypergraph itself is a minimal functional route from *s* to 3.

In fact, by using the incidence matrix of *A* as the stoichiometric matrix, we see that there are no stoichiometric factories from *s* to 3.

We remark that this is not surprising. Computations of elementary flux modes is based on a steady state assumption, that “concentrations” of internal nodes do not change. This is the same assumption for the computation of T-invariants in a Petri Net, where a T-invariant accounts for a preservation of tokens. However, conservation laws for signaling networks are difficult to define due to signal amplification motifs.

## 5 Conclusion

Several different methodologies exist for modeling signaling networks. The reality is that based on the particular components of the signaling network to be studied and the data available, the formalism chosen should be carefully considered, and some-times hybrid approaches are needed [9]. Particularly useful methodologies, when we don’t have enough detailed mechanistic parameters such as kinetic rates of change are graphs, Boolean Networks, hypergraphs and Petri Nets. We have discussed some of the advantages and disadvantages of the different frameworks. Many of the different methodologies for studying signaling networks, *e*.*g*. elementary modes, minimal T-invariants, minimal P-invariants, etc. were introduced for studying reaction networks with a steady state assumption. However, they have successfully been used to capture signaling pathways in mathematical models of signaling networks. The discussion found in [8] for applying elementary flux mode analysis to enzyme cascades show a class of signaling networks where the methodology of T-invariant analysis is directly applicable and not just a formal approach.

We discussed the different methodologies used to estimate how signals are transduced from input layer to target layer. We provided toy examples showing, that although similar, these different methodologies can be strikingly different. Therefore, it is necessary to not only consider the modelling framework, but the appropriate formalism capturing signal transduction. Using only one formalism misses the diversity of strategies the cell uses to transduce a signal.

Some work has been done showing the use of relating the frameworks to each other, which we have discussed in this review [17, 94, 86, 102]. We also related the topological factories to minimal functional routes. It seems reasonable to adopt the concept of topological factories to signaling networks, generalizing the concept of a minimal functional route by allowing nodes to be internally activated, rather than being forced to be activated from an external source. The use of topological factories in signaling networks should be carefully assessed in future work. Relating frameworks with each other opens up the tool-box to analyze how a signal transduces within a cell. Furthermore, understanding how the different methodologies relate to each other will lead to a better understanding of what actually happens *in vivo* inside of a cell.

In graphs with dependent edges, topological factories generalize minimal functional routes. Categorizing graphs with dependent edges such that the concept of minimal functional routes and minimal topological factories are equivalent is of interest. Furthermore, due to the connection between elementary modes with the steady state assumption using the incidence matrix of a graph and simple paths in graphs with no dependent edges [45], it would be interesting to categorize graphs where the minimal S-factories and T-factories are the same.

### 5.1 Author’s contributions

Vera-Licona conceived the study. Sordo Vieira conducted the review and the comparisons under the guidance and supervision of Vera-Licona. Both authors contributed to the writing of the manuscript and approved it prior to submission.

### 5.2 Ethics approval and consent to participate

Not applicable.

### 5.3 Consent to publish

Not applicable.

### 5.4 Availability of data and materials

Not applicable.

### 5.6 Funding

Not applicable.

### 5.7 Graphics

Figure 1 was generated using GINsim, and converted to Encapsulated Postscript (eps) using inkscape. All other Figures were generated using the yEd Graph Editor by yWorks and converted to eps using inkscape.

## 6.1 Abbreviations

MPS: Minimal Path Set
EMT: Epithelial to Mesenchymal transition
MFR: Minimal Functional Route
ESM: Elementary Signaling Mode
S-factory: Stoichiometric factory
T-factory: Topological factory
SF (TF): Stoichiometric (Topological) factory
MSF (MTF): Minimal SF (TF)
T-invariant: Transition invariant
P-invariant: Place invariant
scc: Strongly Connected Component
EGFR: Epidermal Growth Factor Receptor
TGFBR: Transforming Growth Factor Beta Receptor I
FGFR3: Fibroblat Growth Factor Receptor 3
MDM2: Mouse double minute 2 homolog

## 6 Appendix

### 6.2 Concepts from metabolic network analysis

Let 𝒳 be the set of nodes in the source layer, and let *O* be any set of nodes disjoint from 𝒳.

#### Definition 5

*(Topological factory [5, 1]) Given a hypergraph* 𝒢 = (𝒱, ℰ), *a* topological factory *(TF) from X* ⊆ 𝒳 *to O* ⊆ 𝒪 *is a subset* ℱ ⊆ ℰ *with the property that O* ⋃_*e* ∈ ℱ_ *T* (*e*) ⊆ ⋃_*e* ∈ ℱ_ *H*(*e*) ⋃ *X. It is a* minimal topological factory *(MTF) if it contains no smaller topological factory from X to O*.

#### Definition 6

*(Stoichiometric factory [5]) A stoichiometric factory (S-factory) from X* ⊆ 𝒳 *to O ⊆ 𝒪, is a set F ⊆ ℰ such that there exists a flux vector v* ≥ 0 *satisfying:*

1. 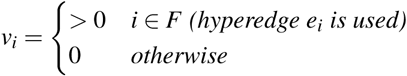

2. (*Sv*)_*V* −*X*_ *≥*0

*3*. (*Sv*)_*O*_ *>* 0

*An S-factory from X to O is minimal* (*MSF*) *if it does not contain any other S-factory from X to O*.

*To include the steady state assumption, we can replace the* ≥ *with* = *in the second constraint for nodes that are neither in X nor O*.

The relationship between topological factories and stoichiometric factories was given in Theorem 1, [5]. We first need to introduce some preliminary concepts to discuss the Theorem and the meaning of it.

#### Definition 7

*(Many-To-One transformation [5]) Let* 𝒢 = (𝒱,ℰ) *be a directed hypergraph. The many-to-one transformation of* 𝒢 *is the directed hypergraph* Φ (𝒢) = (𝒱, Φ (ℰ)) *such that for every e* ∈ ℰ *and for every a* ∈ *H*(*e*), *there is a hyperedge e*_*a*_ ∈ Φ (ℰ) *such that T* (*e*_*a*_) = *T* (*e*) *and H*(*e*_*a*_) = {*a*}.

Given *e* ∈ ℰ, let Φ (*e*) be the set of hyperedges in Φ (ℰ) corresponding to the many-to-one transformation of *e*. If ℱ*⊆* ℰ, let Φ (ℱ) = _*f* ∈ ℱ_Φ (*f*).

Notice that if | *H*(*e*)| = 1, then Φ (*e*) = *e*.

**Observation 1** *If* 𝒢 *is a B-hypergraph, then* Φ (𝒢) =𝒢.

#### Theorem 3

*([5]) For any minimal S-factory H ⊆* ℰ *from X to O in* 𝒢 *there exists a set of minimal topological factories F*_1_,…, *F*_*k*_ *from X to O in* Φ (𝒢) *such that:*

*1. F*_1_, …, *F*_*k*_ *⊆* Φ (*H*);

*2. For each hyperedge r in H, there is i* ∈ {1,∈, *k*} *such that* Φ (*r*) ∩*F*_*i*_ ≠ ∅.

This means that S-factories are the union of minimal topological factories in the many-to-one network. In the case of a B-hypergraph, we immediately get that every minimal S-factory is a union of T-factories.

### 6.3 Elementary Flux Modes

We follow the formalism found in [82] where the mathematical definition of an elementary flux mode was introduced. This has been highly successful for analyzing metabolic networks.

#### Definition 8

*Let* ***N*** *be a matrix in* ℝ^*m×n*^.

*An abstract flux mode* **M** *is the set*

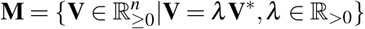

*where* **V**^*\**^ *is a nonzero vector satisfying* **NV**^*\**^ = **0**.

*If* **M** *is a mode with representative* **V**^*\**^, *it is said to be elementary if* **V**^*\**^ *cannot be written as a nontrivial linear combination*

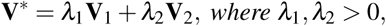

*of nonzero vectors where* 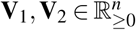 *satisfy* **NV**_*i*_ = **0**, *such that both* **V**_1_, **V**_2_ *contain at least the same number of zero elements as* **V*#x002A** *and at least one of them contains* ; *more zero elements than* **V**^*\**^.

### 6.4 A remark on disjunctive normal form

The definition of disjunctive normal form differs across fields and authors. For example, the definition of disjunctive normal form given in [74] is as follows:

#### Definition 9

*A Boolean function p*(*x*_1_,…, *x*_*n*_) *is said to be in disjunctive normal form if it is the disjunction (connected by the OR operator) of a finite number of terms, each of which has the form e*_1_ ∧*e*_2_ …∧*e*_*n*_ *where e*_*i*_ = *x*_*i*_ *or e*_*i*_ = ¬*x*_*i*_.

In [2], disjunctive normal form simply means that AND clauses are separated by ORs. Thus, (*A* ∧ *B*) ∨ (*C* ∧ *D*) is in disjunctive normal form according to the definition given in [2], whereas it is not in disjunctive normal form according to definition given in [74] since the conjuncts do not contain all variables or their respective negations.

Perhaps more concerning is the following example of two update functions which have the same truth table but are written differently, which can be found in [68]:

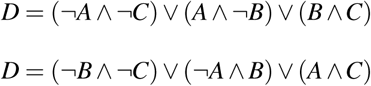

Methods that rely on disjunctive normal form, such as the graph expansion method of [97, 98], might yield different results. One possible solution to this is to write in the so called canonical sum of products form and apply *the same* deterministic logic minimization strategy for reducing the terms (see *i*.*e*. [86] for references to logic minimization algorithms).

